# Dystonia-specific mutations in *THAP1* alter transcription of genes associated with neurodevelopment and myelin

**DOI:** 10.1101/2021.06.22.449452

**Authors:** Aloysius Domingo, Rachita Yadav, Shivangi Shah, William T. Hendriks, Serkan Erdin, Dadi Gao, Kathryn O’Keefe, Benjamin Currall, James F. Gusella, Nutan Sharma, Laurie J. Ozelius, Michelle E. Ehrlich, Michael E. Talkowski, D. Cristopher Bragg

## Abstract

Dystonia is a neurologic disorder associated with an increasingly large number of variants in many genes, resulting in characteristic disturbances in volitional movement. Dissecting the relationships between these mutations and their functional outcomes is a critical step in understanding the key pathways that drive dystonia pathogenesis. Here we established a pipeline for characterizing an allelic series of dystonia-specific mutations in isogenic induced pluripotent stem cells (iPSCs). We used this strategy to investigate the molecular consequences of variation in *THAP1*, which encodes a transcription factor that has been linked to neural differentiation. Multiple pathogenic mutations that have been associated with dystonia cluster within distinct THAP1 functional domains and are predicted to alter its DNA binding properties and/or protein interactions differently, yet the relative impact of these varied changes on molecular signatures and neural deficits is unclear. To determine the effects of these mutations on THAP1 transcriptional activity, we engineered an allelic series of eight mutations in a common iPSC background and differentiated these lines into a panel of near-isogenic neural stem cells (n = 94 lines). Transcriptome profiling of these neural derivatives followed by joint analysis of the most robust individual signatures across mutations identified a convergent pattern of dysregulated genes functionally related to neurodevelopment, lysosomal lipid metabolism, and myelin. Based on these observations, we examined mice bearing *Thap1*-disruptive alleles and detected significant changes in myelin gene expression and reduction of myelin structural integrity relative to tissue from control mice. These results suggest that deficits in neurodevelopment and myelination are common consequences of dystonia-associated *THAP1* mutations and highlight the potential role of neuron-glial interactions in the pathogenesis of dystonia.

## INTRODUCTION

The dystonias are a group of complex neurologic disorders characterized by sustained involuntary muscle contractions and twisted postures.^1^ A multitude of pathogenic mechanisms has been proposed, most of which relate broadly to signaling cascades within neurons.^2–4^ A small number of genes have been unambiguously associated with specific phenotypes,^5,6^ but the existence of shared molecular pathways disrupted across dystonic syndromes remains uncertain. Moreover, in several genes associated with dystonia, the functional implications of diverse genetic perturbations (e.g., loss-of-function and missense point mutations or copy number variation) remain largely unexplored. Establishing the neurobiological basis for dystonia across the diverse genes and mutations underlying its genetic architecture can provide insights into other neurodegenerative and/or neuropsychiatric disorders, given the many anatomic, neurophysiologic, and even phenotypic overlaps in these diseases, raising the potential for broadly applicable therapeutic intervention.

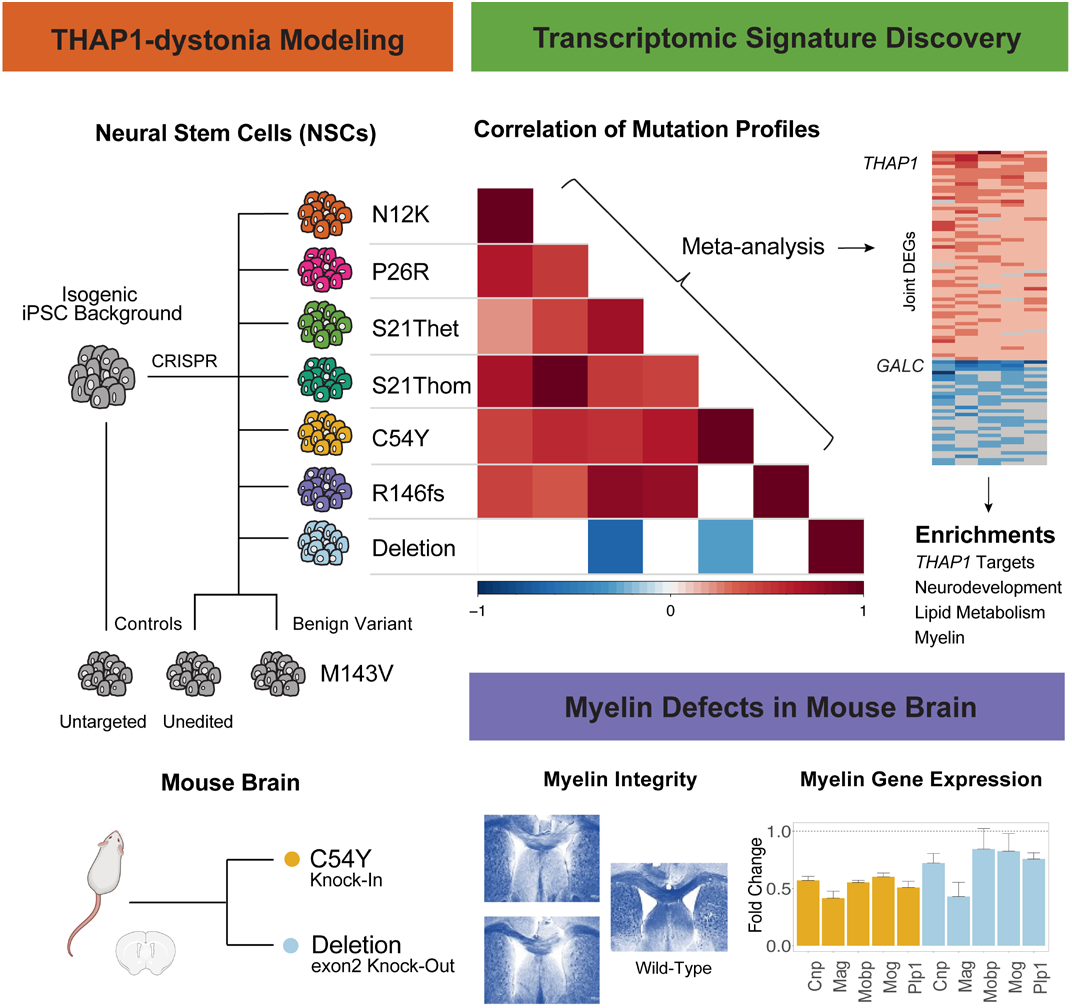

The integration of induced pluripotent stem cell (iPSC) modeling and genome editing technologies offers an emerging opportunity to study mutational mechanisms in cells specifically relevant to disorders of the nervous system.^7^ Combining these cellular modeling tools with transcriptomic profiling can be a powerful approach to identify shared molecular signatures that converge on common affected pathways, as well as molecular changes associated with distinct mutations.^8–10^ However, there are considerable technical and biological barriers to such an approach, and there have been few examples of CRISPR modeling of disease-associated mutational mechanisms in near-isogenic neuronal models of disease.^11^ Here we applied this strategy to explore the functional impact of genetic variation in *THAP1* (Thanatos-associated [THAP]-domain containing Apoptosis-associated Protein 1), which causes familial and sporadic cases of dystonia.^12^

The THAP1 protein is a zinc finger transcription factor consisting of an N-terminal DNA-binding domain, a central proline-rich region, and a C-terminal coiled-coil domain bearing a leucine zipper motif and nuclear localization signal (NLS).^13–15^ The cellular functions regulated by THAP1 are not fully defined, but studies in mice have linked it to the reciprocal repression of the pluripotency program and the promotion of neural differentiation via activation of progenitor-specific genes in embryonic stem cells.^16^ Missense and truncating variants in the DNA-binding domain represent the majority of known pathogenic variants, but biophysical studies suggest that they may affect the protein in different ways by: (1) altering affinity for the THAP1 DNA-binding sequence; (2) changing the specificity for this element; or (3) decreasing protein stability.^17,18^

Less is known about variation within C-terminal domains, although ones that alter key residues within the NLS and leucine zipper have been suggested to interfere with nuclear translocation and homodimerization, respectively.^15,19,20^ Most of these variants occur exclusively in dystonia probands, although some are found in large control databases, suggesting that they represent either dystonia-associated mutations that display reduced penetrance or benign variation.

Here we engineered an allelic series of mutations in *THAP1* and performed transcriptome analyses to specifically interrogate varied mutational changes and consequent functional alterations in neural stem cells (NSCs). The NSCs represent multipotent cells within the germinal zone that precede specification into neuronal and glial lineages and give rise to neurons, oligodendrocytes, and astrocytes within the adult brain.^21,22^ Transcriptional profiling identified a core set of dysregulated genes in NSCs across diverse *THAP1* mutations that converge on signatures related to neurodevelopment and myelination. We further assessed myelin status directly in mice with *Thap1* disruption and confirmed that these perturbed pathways lead to dysregulated myelin gene expression and reduced myelin structural integrity in adult animals. Our results underscore previously unappreciated neurodevelopmental aspects of *THAP1*-dystonia and highlight the potential contribution of glial dysfunction to mechanisms of neurologic diseases.

## MATERIALS AND METHODS

### CRISPR Engineering of THAP1 Variants, NSC Differentiation

Single-guide RNAs targeting *THAP1* mutations were designed using the known genomic sequence of *THAP1* (GRCh37, ENSG00000131931) and the Benchling CRISPR tool, and synthesized by Synthego. To generate the genomic deletion, double-guide RNAs were used; these were designed using sequences in the 5, and 3, untranslated regions. A karyotypically normal human male iPSC line from an individual without known disease (33362D, passage <30) was seeded at 100,000 cells per well and maintained on mTESR1 media (Stemcell Technologies). On day three post-plating and upon reaching 50% confluence, colonies were pretreated with nocodazole 25ng/uL media overnight, then transfected the next day with Lipofectamine Stem (Thermofisher), the appropriate guide-RNA, Cas9 protein (Invitrogen), a single-stranded 100-bp oligo donor nucleotide (ssODN) template, and GFP-mRNA (TriLink Biotechnologies). Fluorescence was used as a proxy of transfection efficiency and for sorting positive cells at day three post-transfection. GFP-positive, sorted cells were divided into ‘sibs’, seeded into 16 wells of a 96-well plate, and maintained until confluent.^23^ At this point, colonies were still a mixture of edited and unedited iPSCs. Editing efficiency within each sib was assessed using digital droplet PCR (ddPCR) and Genome Edit Detection assays (Bio-Rad). Two sibs with highest editing efficiencies were selected, divided into 16 sibs, replated and regrown. This sib-selection method was repeated until edit efficiency within a sib reached >5% or within three rounds of assaying. Eventually, a selected sib was expanded, then sorted as single-cells into wells of a 96-well plate. DNA was extracted and Sanger sequencing was performed to identify clonal populations bearing desired mutations in *THAP1*. Independent mutation-positive lines and unedited controls were expanded (average = four per mutation). Prior to cryopreservation for future differentiation, another round of Sanger sequencing was performed to verify the mutation and clonality, and up to four random clones from each group were subjected to routine karyotyping (WiCell), which did not reveal any abnormalities. To identify and verify *THAP1*-deletion cell lines, a combination of PCR and fragment length analysis on agarose gels, Sanger sequencing, and ddPCR-based copy number assays (Biorad) was used. Guide RNAs, oligonucleotide templates, and ddPCR/Sanger sequencing primers are available from the authors upon request.

Cryopreserved iPSCs were thawed and grown on mTESR1 media for three passages prior to induction of neural differentiation. Differentiation into NSCs was performed following standard manufacturer’s protocol (Gibco). Briefly, this involves a seven-day neural induction stage until passage zero, followed by expansion. All inductions were performed using supplements from the same batch/lot, and all NSCs used for further experiments were from passage three. Importantly, differentiation batches followed analysis batches. Within the same CRISPR experimental groups, mutation-positive/edited and unedited samples were differentiated together, with equal numbers of mutant and control samples split into two differentiation batches per group. This resulted in eight differentiation batches for the entire study, with each batch containing balanced numbers of mutant and control samples from two different CRISPR experiments. To assess efficiency of neural differentiation, up to four randomly selected samples per differentiation batch were subjected to immunocytochemical staining with known markers of NSCs (Nestin, Msi1, Pax6) (***Supplementary Figure 1***). The pluripotency marker Oct4 was negative in all samples tested (data not shown).

### Library Preparation and RNA Sequencing, Quality Control

Once all differentiations were complete, RNA was extracted from all samples using Trizol lysis and a standard RNA isolation kit (Qiagen). One microgram of each RNA sample was taken for library preparation using the Illumina TruSeq protocol and adapters. One-hundred-and-fifty-base pair, paired-end reads were generated using a NovaSeq S4 (Broad Institute Genomics Platform). Up to 65 million reads (mean: 38.4 million, median: 37.8 million) were generated per sample, resulting in a median of 73.2× mean per base coverage per sample. Sequence adapters were trimmed using Trimmomatic, then aligned to the Homo sapiens GRCh37.75 reference sequence using the STAR Aligner (version 2.5.3).^24^ As part of post-alignment quality control, two samples with > 7% intergenic rate were excluded from downstream analysis. Further, mutation loci were verified by visualizing using the Integrative Genomics Viewer (IGV version 2.3.94). From this, one sample was relabeled from N12K_5 (mutant) to N12wt_6 (control). Counts were generated from the STAR Aligner. Allele specific expression was performed using the Genome Analysis ToolKit (GATK version 3.5) ASEReadCounter tool, and a binomial test was applied to determine significant deviation from the expected equal expression of reference and variant alleles (p-value < 0.05). Off-target effects secondary to CRISPR editing were investigated using CasOFF-Finder.^25^ Exonic off-target hits with up to three mismatches from the on-target site were queried via samtools for the presence of alternative alleles. Among sites investigated, a missense variant in Chr3:119886818 (GPR1) was seen in two mutant (P26R_4, P26R_6) and two control lines (P26wt_3, P26wt_4); otherwise, no other relevant exonic off-target effects were identified in other groups.

### Differential Expression Analyses, Meta-Analysis

Differential expression analysis was performed using the DESeq2 package for R (version 1.26.0).^26^ Analysis was performed for each mutation individually using mutation-positive samples and control samples within the same NSC differentiation batches (n = 8). Our batching protocol allowed us to exploit controls from parallel genome editing experiments to increase sample size in controls while keeping analyses within batches. Thus, there were four to six mutant profiles and 11 to 13 internal control profiles per comparison in each optimized differential gene expression experiment. Prior to performing DESeq2 analysis, data was filtered by including only genes with > 0.5 counts per million (cpm) in >= 60% of samples within an analysis group, resulting in an average of 16493 ‘expressed genes’ analyzed per group, then normalized using DESEq2’s median of ratios method.^26^ Exploratory principal component analysis (PCA) using uncorrected expression profiles revealed that differentiation batch effects dominated the variance. Visual comparison of PCAs revealed that profiles appropriately clustered according to genotype after application of surrogate variable (SV) analysis using the sva package (version 3.34.0),^27^ and correcting for the SVs in the regression models (***Supplementary Figure 2***). Differentially expressed genes (DEGs) were thus obtained from SV-corrected counts and using a predefined cutoff of false-discovery rate (FDR)-adjusted p-value < 0.05. Fold-change (FC) was estimated from DESeq2, and a cutoff of log2FC > 0.59 or log2FC < −0.59 was used to determine significant DEGs. To express log2FCs to FC, we used x = 2^(log2FC). Annotation DB (version 1.48.0) was used to obtain gene names from Ensembl IDs. To generate models for overlapping DEGs, we used background expectation rates using the SuperExact R package.^28^ Fisher’s exact test p-values were then adjusted using the Bonferroni method according to the number of overall set intersection probabilities.

To compare expression profiles in different comparisons beyond overlapping DEGs, rank-scores per gene (q-values) were derived from signed -(log10) transformed p-values in each comparison. Spearman’s pairwise rank correlation was then obtained for every comparison group, using different thresholds to subset q-values and obtain correlation patterns. Subsequently, a meta-analytic approach was applied to obtain a core set of DEGs across multiple mutations in the DNA-binding domain and NLS. First, per-gene two-sample means Z-score testing was performed between mutant and control SV-corrected counts. These scores were then weighted according to the total number of samples within that comparison.^29^ Meta-analysis Z-scores per gene were obtained using weighted Z-scores from the N12K, S21Thet, P26R, C54Y, and R146fs comparison groups. From these, _JOINT_DEGs were determined using a cutoff of Benjamini-Hochberg (BH)-adjusted meta-analysis p-value < 0.01 and a sign-test, which required all individual components to be in the same direction of expression change.

### Functional Enrichment Analyses, Weighted Gene Coexpression Network Analysis

Gene Ontology (GO) term enrichment analysis was performed using _JOINT_DEGs and _DEL_DEGs (FDR < 0.01), using the companion R package for g:Profiler2 (version 0.2.0).^30^ To reduce these lists for visualization, semantic similarity was applied using the SimRel measure in REVIGO to generate a small list (allowed similarity 0.5) of related significantly enriched GO terms.^31,32^ Reduced lists (from _DEL_DEGs) were additionally manually curated to show common enriched GO terms in up- and downregulated DEGs. Consensus THAP1 targets were obtained from the gene set M30208 (‘THAP1_TARGET_GENES’) in the MSigDB database (https://www.gsea-msigdb.org/), which derived results from ChIP-Seq experiments uploaded into the Gene Transcription Regulation Database.^33^ A hypergeometric test was applied to determine significant overlap with _JOINT_DEGs and _DEL_DEGs.

Coexpression analysis was performed using the WGCNA R package (version 1.69).^34^ Expression count matrices (n = 87 samples after removal of outliers) were subsetted to include only protein-coding genes (total genes = 12661 genes), then used to construct signed networks from adjacency and topological overlap matrices at soft power threshold of 15, which was calculated to reduce mean connectivity < 100, as suggested by package authors. This resulted in 59 modules of correlated genes, then reduced to 47 after merging with the mergeCloseModules function. An additional custom filtering step was then applied wherein only genes with module membership p-value of < 0.10 were considered to belong to their assigned modules; otherwise, genes were assigned to M0 (‘grey’/ unclassified module, n = 702 genes). Modules of interest were identified by obtaining the biweight mid-correlation between the module eigengene (ME) and *THAP1* expression,^35^ and by testing for significant enrichment of _JOINT_DEGs and _DEL_DEGs in each module, using a Fisher’s test (p-value < 0.10). To derive functional insights from these modules of genes, GO term enrichment analysis was performed as above. MSigDB was likewise queried for significant overlap with hallmark non-cancer related gene sets (version 7.2, build date September 2020). Additionally, significant overrepresentation of module genes in custom lists was tested. These lists were assembled based on hypotheses derived from known functions of *THAP1* as well as enriched GO terms in modules of interest, and included: a list of genes related to lysosomal regulatory pathways,^36^ a list of human myelin-related genes,^37^ polycomb genes related to developmental regulators in human stem cell populations,^38^ and modules of genes in different periods of brain development derived from functional genomic analyses (PsychEncode data set).^39^ Overrepresentation in these and other brain-related gene lists was investigated using Fisher’s tests, applied through the WGCNA and anRichment R packages (ver 1.18), and adjusted for multiple testing. For all enrichment testing (GO/custom lists/ DEGs), custom background sets corresponding to genes assayed were used (‘expressed genes’, mean 16303 genes across all comparisons), or to protein-coding genes (12661 genes, for enrichment testing in WGCNA-derived results).

### Genetic Mouse Models, Myelin Expression and Staining

Experiments in mice were in compliance with the United States Public Health Service’s Policy on Humane Care and Use of Experimental Animals and were approved by the Institutional Animal Care and Use Committee at Icahn School of Medicine at Mount Sinai. Genetic mouse models included heterozygote *Thap1*^C54Y/+^ (C54Y-KI) and *Thap1*^-/+^ (ex2-KO) mice, backcrossed over 10 times onto a C57Bl6/J background. Controls were wild-type (WT) littermates of each mutant genotype. For targeted gene expression assays of myelin-specific genes, tissue from the cortex, cerebellum, and striatum of postnatal day 21 (P21, n = 5 C54Y-KI, 7 WT littermates; n = 10 ex2-KO, 12 WT littermates) and two-month-old mice (adult, n = 4 per group) were lysed and homogenized in QIAzol Lysis Reagent (Qiagen). Total RNA purification was performed with the miRNeasy Micro Kit (Qiagen) according to manufacturer’s instructions. 500 ng of RNA were reverse-transcribed using the High Capacity RNA-to-cDNA Kit (Applied Biosystems). Real-time quantitative PCR was then performed on resulting cDNAs in a Step-One Plus system (Applied Biosystems) using PerfeCTa SYBR Green FastMix ROX (Quanta BioSciences). Quantitative PCR for each target consisted of 40 cycles, each with 15 seconds at 95°C and 30 seconds at 60°C, followed by dissociation curve analysis. Primer oligonucleotide sequences are available upon request. Gene expression levels were obtained using the ΔΔCt method (normalized to *Gapdh* expression), then expressed as mean FC relative to the mean expression in the control group.

For immunocytochemical staining of myelin-specific genes, mice (adult, n = 4 C54Y-KI, 5 ex2-KO, WT = 7 mice) were anesthetized in a CO_2_ chamber and transcardially perfused with ice-cold phosphate buffer solution (PBS). Brains were post-fixed in 4% polymeric formaldehyde and sliced into 30 μm sections using a vibratome (Leica) for histological analyses. Sections were permeabilized with 0.1% Triton X-100 in PBS, blocked with PBS/5% goat serum, and then incubated with primary antibodies: Mobp (1/1000, rabbit, Abgent) and Cnp (1/1000 mouse, Abcam) prior to exposure to fluorescent Alexa secondary antibodies. Immunoreactive proteins were visualized using Enhanced Chemiluminescence on a Fujifilm LAS4000 imaging device, and optical densities of protein of interest were measured and analyzed with Fiji software (ImageJ). For non-fluorescent myelin immunostaining, luxol fast blue (LFB) stain was used to measure myelin structural integrity in the anterior commissure and corpus callosum of adult mice (n = 4 C54Y-KI, 5 WT littermates; n = 8 ex2-KO, 8 WT littermates). Briefly, sections were dried for 30 minutes before being immersed in a de-fat solution (1:1 alcohol/chloroform for 2h), after which, sections were transferred to 0.1% LFB overnight at 56°C. After several washes in 95% ethyl alcohol, followed by distilled water, the sections were first differentiated in a lithium carbonate solution for 30 seconds, followed by a second differentiation in 70% ethyl alcohol for 30 seconds. Sections were checked microscopically to verify the sharp definition of gray versus white matter before being dehydrated in graded alcohol baths, cleared in xylene, and mounted with resinous medium. Images were obtained with an Olympus BX61 microscope and analyzed with Fiji software (ImageJ). Quantification was performed and Student’s t-tests were used to determine significance in difference between quantification in mutant and WT littermates (in the anterior commissure measurements were taken from both left and right).

## RESULTS

### Generation of an allelic series of THAP1 mutations in neural stem cells

To define the transcriptional changes of *THAP1*-associated mutational mechanisms, we focused primarily on mutations within the DNA-binding domain given that: (1) 70% of pathogenic sequence variation associated with *THAP1*-dystonia are found in this module,^40^ and (2) mutations in this region have been proposed to directly alter the protein’s transcriptional activity^14,18^ (***Figure 1A***). We chose variants based on their distribution within the three-dimensional structure of this domain, which is composed of an anti-parallel β-sheet and a helix-loop-helix motif.^18,41^ This allelic series consisted of four missense mutations: N12K and S21T, which are localized within the first loop (L1) of this domain; P26R within the second loop (L2); and C54Y within the β-sheet.^17^ All four mutations are: (1) documented in dystonia probands^42–44^; (2) absent from population-based genetic references in the Genome Aggregation Database (gnomAD)^45^; (3) predicted to be damaging across multiple algorithms (Combined Annotation Dependent Depletion [CADD], Mendelian Clinically Applicable Pathogenicity [M-CAP], Rare Exome Variant Ensemble Learner [REVEL])^46–48^; and (4) classified as pathogenic based on existing criteria^49^ (***Figure 1A***). For comparison, we also engineered iPSCs bearing two genotypes to model *THAP1* loss-of-function and haploinsufficiency, respectively: (1) a pathogenic dystonia-specific frameshift mutation, R146fs,^50^ that results in a prematurely terminated protein and prevents nuclear translocation by abolishing the NLS; and (2) a deletion allele (DEL), in which the entire coding sequence was excised using a dual guide-RNA approach. To date there has been one report of a family affected with a severe and atypical form of dystonia carrying a segregating ~500-kilobase genomic deletion of the 8p11.21 locus that included all exons of *THAP1*,^51^ and there are no *THAP1* gene deletions in gnomAD-SV v2.1.^52^ Finally, we generated iPSCs bearing a missense mutation, M143V, within the C-terminal coiled-coil domain. Although this variant has been described recurrently in dystonia families,^53–55^ its frequency is relatively high in gnomAD, i.e., above the maximum tolerated allele frequency in a reference database for *THAP1* and for an estimated disease penetrance of 60%.^56^ Thus, we hypothesized that this variant may be benign and would behave as a control.

**Figure 1.**
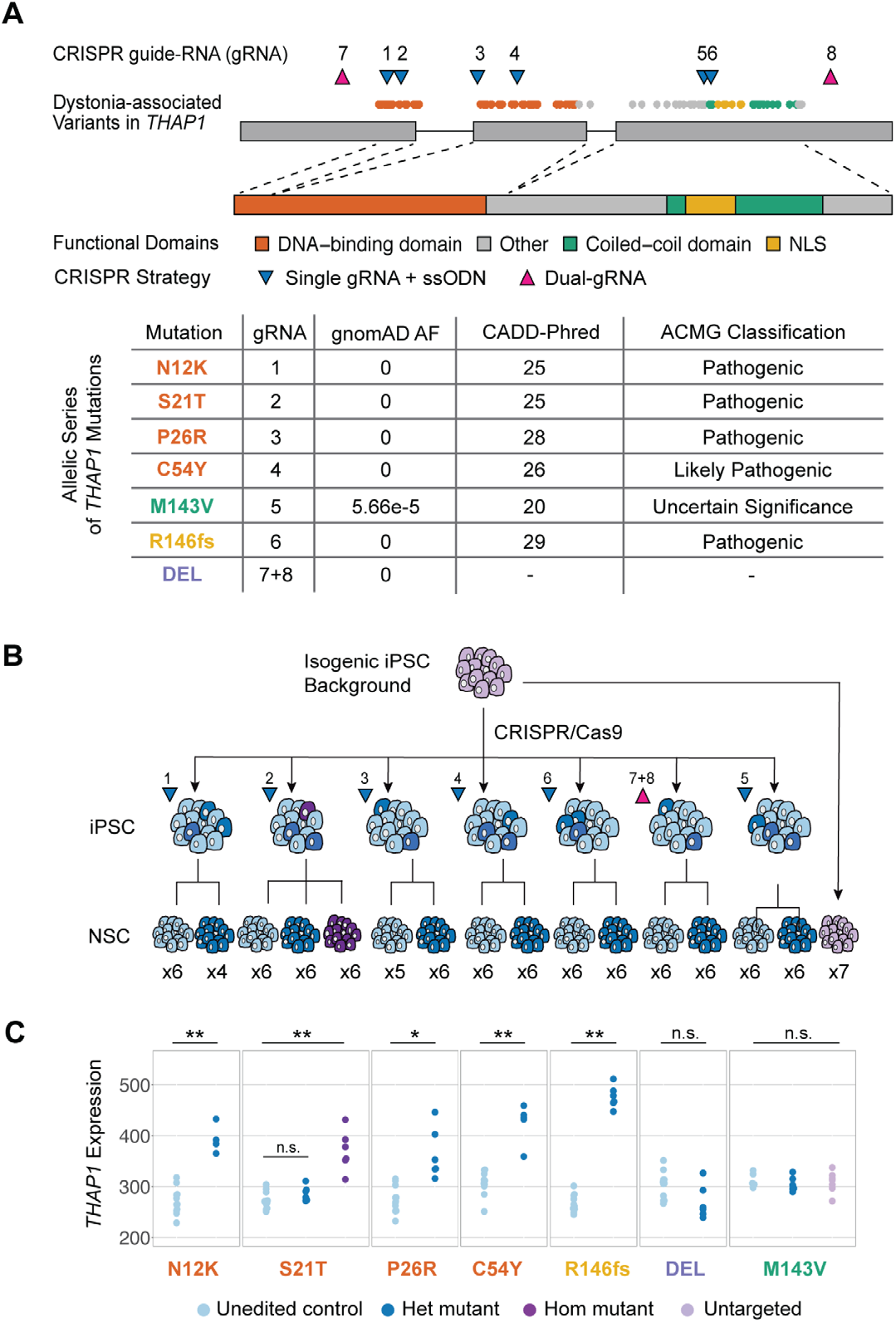
*Generation of an allelic series of* THAP1 *mutations in neural stem cells*. **(A)**Dystonia-associated genetic variation in *THAP1* localize within known functional domains of the protein, with 70% of disease-specific, pathogenic mutations corresponding to residues in the DNA-binding domain. To create an allelic series, four single-nucleotide variants in this domain (N12K, S21T, P26R, C54Y) and one in the coiled-coil domain (M143V) were introduced via CRISPR/Cas9-based genome editing, and using single-guide RNAs (gRNAs) in combination with a single-stranded oligo donor nucleotide (ssODN) template. An eight base-pair small deletion resulting in a frameshift (R146fs) in the NLS was generated using the same approach. Among these, mutations in the DNA-binding domain and the frameshift in the NLS are considered pathogenic based on absence in gnomAD, a large population-based database of exome sequences that is depleted of individuals with neurologic disorders, bioinformatic predictive methods, including the CADD Phred-score, and classification using criteria from the American College of Medical Genetics and Genomics (ACMG). The M143V is classified as a variant of unknown significance using these criteria. Additionally, gRNAs placed in untranslated regions of the gene were used to engineer a haploinsufficiency model via deletion of the entire coding sequence of *THAP1* (DEL). AF column denotes allele frequency in gnomAD v2.1.1. **(B)** Mutations were generated from a single human induced pluripotent stem cell (iPSC) line (“background”), making the models isogenic beyond the mutation introduced in *THAP1*, and effectively isolating mutational effects in subsequent experiments. Multiple independently edited and matched unedited cells from each CRISPR experiment were differentiated into NSCs. For each mutation, up to six edited NSC lines and up to six unedited control lines were used. Additionally, six NSC lines were generated from an iPSC line bearing a homozygous S21T mutation, and seven NSC lines that were directly differentiated from the background line were also used as controls (total n = 94 NSC lines). See also ***Supplementary Figure 1***. **(C)** *THAP1* expression was assessed in each mutation vs. corresponding control comparison. The results recapitulate previously described upregulation of *THAP1* in lines carrying pathogenic genetic variation, i.e., heterozygous (Het) N12K, P26R, C54Y and R146fs mutations, and homozygous (Hom) S21T. The DEL model induced minimal reduction of THAP1 expression, while gene expression was unchanged in M143V NSCs. *THAP1* expression plotted as normalized counts. Wald test statistic from differential expression analysis: * p-value < 0.05; ** FDR < 0.05; n.s. not significant.

The CRISPR targeting method across seven genome editing sites produced individual iPSC clones that were heterozygous for each mutation. For the S21T locus, the editing strategy generated clones with heterozygous (S21Thet) and homozygous (S21Thom) alleles in the same experiment, and we included the latter in transcriptional profiling to investigate potential dosage effects. Multiple independently edited iPSC clones and matched unedited controls that were exposed to the same guide RNAs, as well as controls that were directly isolated from the background, were differentiated into NSCs in predefined batches (four to seven NSC lines per genotype per mutation, total n = 94 NSC lines) (***Figure 1B***). There were no obvious growth or morphologic differences observed between mutant and control iPSC/NSC lines, and equivalent expression of neural markers was seen across differentiation batches (***Supplementary Figure 1***). Based on previous reports of *THAP1* autoregulation,^16,57^ an interesting aspect of these analyses was the opportunity to directly quantify the expression of *THAP1* across all mutations in the allelic series. From these analyses, we were able to note: (1) significant upregulation of *THAP1* when disrupted by a dystonia-specific pathogenic variant (mean FC = 1.4 across lines with a heterozygous N12K, S21T, P26R, C54Y, or R146fs mutation); (2) minimal impact of complete deletion of the *THAP1* coding sequence on expression, suggesting strong autoregulation of *THAP1* (FC = 0.89, p-value 0.08); and (3) no change in *THAP1* expression by M143V (***Figure 1C***). While allele-specific expression was not significantly different between reference and variant alleles in the heterozygous C54Y, P26R, S21T, and M143V groups, expression of the reference allele was moderately increased in the N12K lines (mean fraction of reference allele/ total = 0.72; p-value 0.05, binomial test).

### Pathogenic variation in the *THAP1* DNA-binding domain and nuclear localization signal induce overlapping transcriptome-wide alterations

The mutations we engineered localize to different THAP1 structural and functional domains. We therefore asked if they produced distinct patterns of transcriptional dysregulation in differential expression analyses within experimental groups and by matching mutations to unedited controls within differentiation batches. After controlling for variance in global expression profiles associated with batch effects from editing and differentiation, principal component analyses revealed that all profiles clustered according to genotype (***Supplementary Figure 2***). For each comparison, we identified a set of DEGs (FDR < 0.05, designated hereafter as significant if FC > 1.5). Full gene deletion resulted in the most profound transcriptome-wide perturbation (2622 _DEL_DEGs, 386 significant), followed by introduction of a homozygous missense mutation (533 _S21Thom_DEGs, 130 significant) (***Figure 2A***).

**Figure 2.**
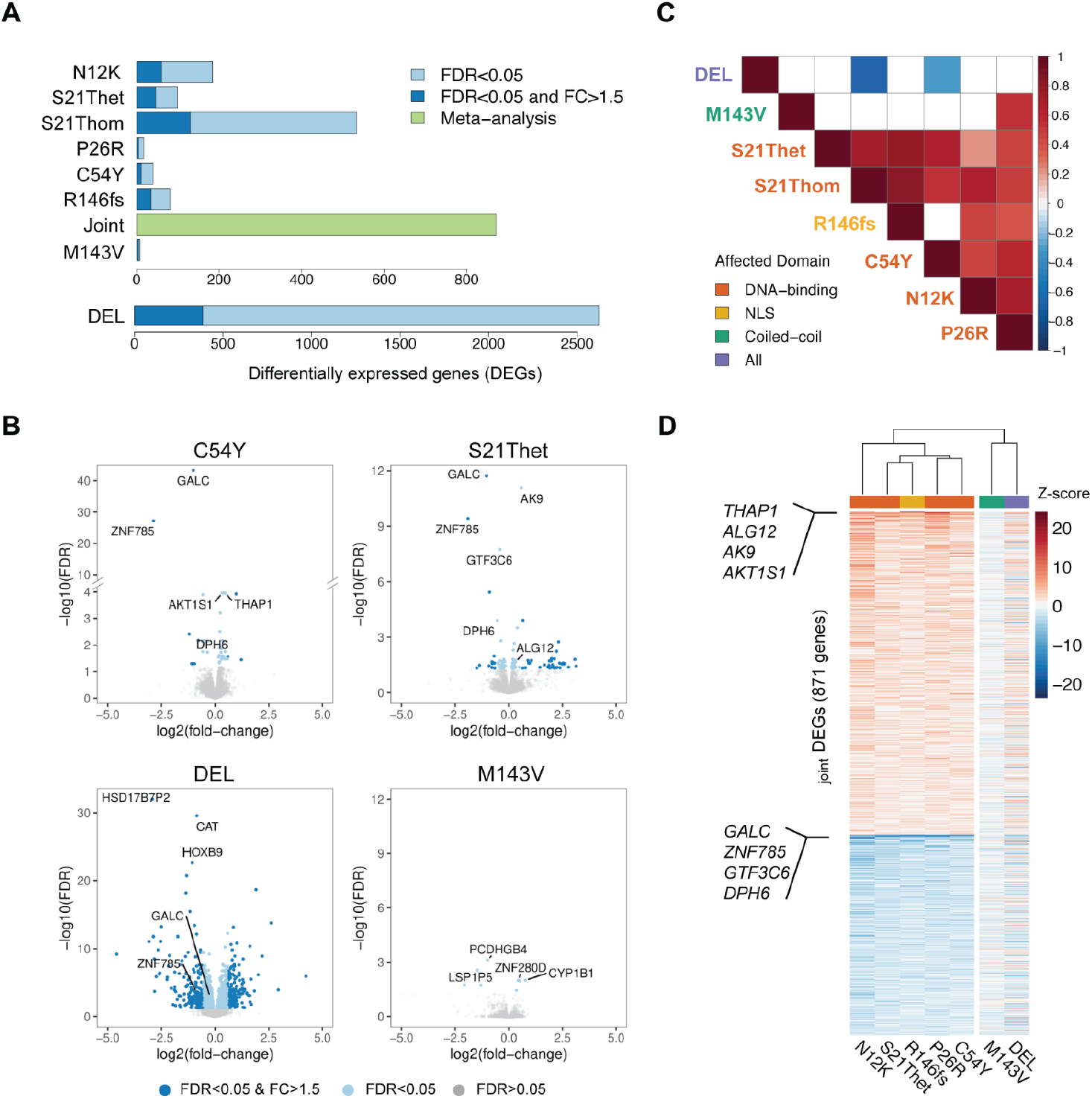
Pathogenic variation in the THAP1 DNA-binding domain and nuclear localization signal induce overlapping transcriptome-wide alterations. **(A)** Differential expression analysis was performed within mutational groups independently, identifying a set of robust and significant DEGs for each comparison (FDR < 0.05, FC > 1.5). Full gene deletion resulted in the most profound transcriptome-wide perturbation (2622 _DEL_DEGS, 386 significant), followed by introduction of a homozygous missense mutation (533 _S21Thom_DEGs, 130 significant). _JOINT_DEGs are genes identified via meta-analytic combination of results in N12K, S21Thet, P26R, C54Y, and R146fs groups. **(B)** Volcano plots showing DEG discovery in selected mutational groups: C54Y and S21Thet (representing DNA-binding domain mutations), DEL, and M143V, with notable overlapping DEGs induced by mutations localized to the DNA-binding domain labeled (i.e., C54Y and S21Thet; upregulation of *THAP1*, *ALG12*, *AK9*, and *AKT1S1*, and downregulation of *GALC*, *ZNF785*, *GTF3C6* and *DPH6*). These were also the most consistently dysregulated genes in the joint analysis. _DEL_DEGs intersected with few DEGs induced by other pathogenic mutations (e.g., *GALC* and *ZNF785*). In contrast, there were only a few DEGs induced by M143V. See also ***Supplementary Figures 2 and 3***. **(C)** Pairwise correlations of extreme rank-orders of genes revealed a striking overlap in the expression signatures induced by missense mutations in the DNA-binding domain and by the frameshift in the NLS (mean Spearman rank correlation r = 0.5, p-values < 0.05; outlier: C54Y-R146fs), consistent with the observation of overlapping DEGs discovered in these mutational groups. See also *Supplementary Figure 4*.

The experimental design of having mutants and controls within differentiation batches facilitated independent differential expression analyses and subsequent enrichment tests for shared DEGs across mutations by comparison to null expectation models.^28^ Intersections of DEGs induced by any two heterozygous mutations localized to either the DNA-binding domain or NLS were always significantly above expectation (minimum 9.6 fold-enrichment of observed/expected overlapping DEGs, Bonferroni-corrected p-value [FWER] 1.1×10^-6^ [_N12K_DEGs-_S21Thet_DEGs]). This observation was consistent even when considering multi-set intersections (i.e., overlaps across 3 to 6 DEG sets), indicating a significant overlap in the transcriptional consequences of mutations affecting these functional domains. The most consistently observed effect was downregulation of *GALC*, *ZNF785*, and *DPH6* across all six mutational groups (3.7×10^9^ fold-enrichment over expected, FWER 1.6×10^-26^), and dysregulation of dosage-sensitive DEGs induced by S21Thet/hom (7.7-fold-enrichment of observed/expected overlapping DEGs, FWER 2.4×10^-15^), where 78% of overlapping DEGs displayed greater fold-changes associated with the homozygous alteration (***Supplementary Figure 3***). Importantly, while only 0.6% of _DEL_DEGs intersected with DEGs induced by any mutation localized to either the DNA-binding domain or NLS, closer inspection of expression profiles indicated consistent expression changes in the DEL group at nominal thresholds (***Figure 2B*** and ***Supplementary Figure 3***). These results suggest that pathogenic *THAP1* mutations affecting the DNA-binding domain and NLS exert highly concordant transcriptomic effects and that while these molecular signatures were largely distinct from those observed in haploinsufficiency, there were potentially convergent signatures across all mutational mechanisms.

We further explored the common transcriptional signatures using pairwise correlations of extreme rank-orders of genes (i.e., upper and lower 2.5% of q-values, corresponding to up to the 830 most differential genes in every set). In this subset of expression profiles, the rank-orders in DNA-binding domain mutants correlated with each other and with those in the NLS group (mean Spearman r = 0.56; p-value < 0.05; outlier: C54Y-R146fs), but not with DEL or M143V q-values (***Figure 2C***). Further, the correlation patterns were maintained at different thresholds (***Supplementary Figure 4***), suggesting a robust intersecting signature induced by the N12K, S21T, P26R, C54Y, and R146fs mutations that extend beyond the few overlapping significant DEGs already discovered. Meta-analysis using signed weighted Z-scores further defined a molecular signature consisting of 871 genes (termed subsequently as _JOINT_DEGs, representing meta-analysis FDR < 0.01) that were consistently altered in a direction-specific manner across these dystonia-specific pathogenic mutations in the DNA-binding domain and the NLS (***Figure 2D***).

### Functional analysis reveals convergent signatures of neurodevelopment and myelination across diverse *THAP1* mutations

To identify potential cellular consequences of *THAP1* transcriptional dysregulation, we performed functional enrichment analyses of the discovered DEGs: (1) genes identified via meta-analysis (_JOINT_DEGs), representing the common signature of genetic variation in the DNA-binding domain and NLS, and (2) _DEL_DEGs, representing haploinsufficiency. The analyses revealed significant enrichment of multiple GO terms primarily related to transcription regulation in _JOINT_DEGs (FDR < 0.05, Fisher’s test, ***Figure 3A***), consistent with the characterization of THAP1 as a transcription factor.^14,58^ Further, _JOINT_DEGs overlapped significantly with consensus THAP1 target genes^33^ as expected (FDR 1.8×10^-9^), given that the mutations that comprise this aggregation either functionally impact THAP1 DNA-binding or impair nuclear import.^14,17,19^ Meanwhile, _DEL_DEGs were strongly related to developmental processes including nervous system development (FDR 2.3×10^-7^) (***Figure 3B***), recapitulating previous observations in *Thap1*-haploinsufficient mouse embryonic stem cells.^16,59^ Intriguingly, enrichment analysis in _DEL_DEGs also revealed enrichment of terms related to lipid metabolism and synthesis (e.g., steroid biosynthetic process, FDR 3.4×10^-5^).

**Figure 3.**
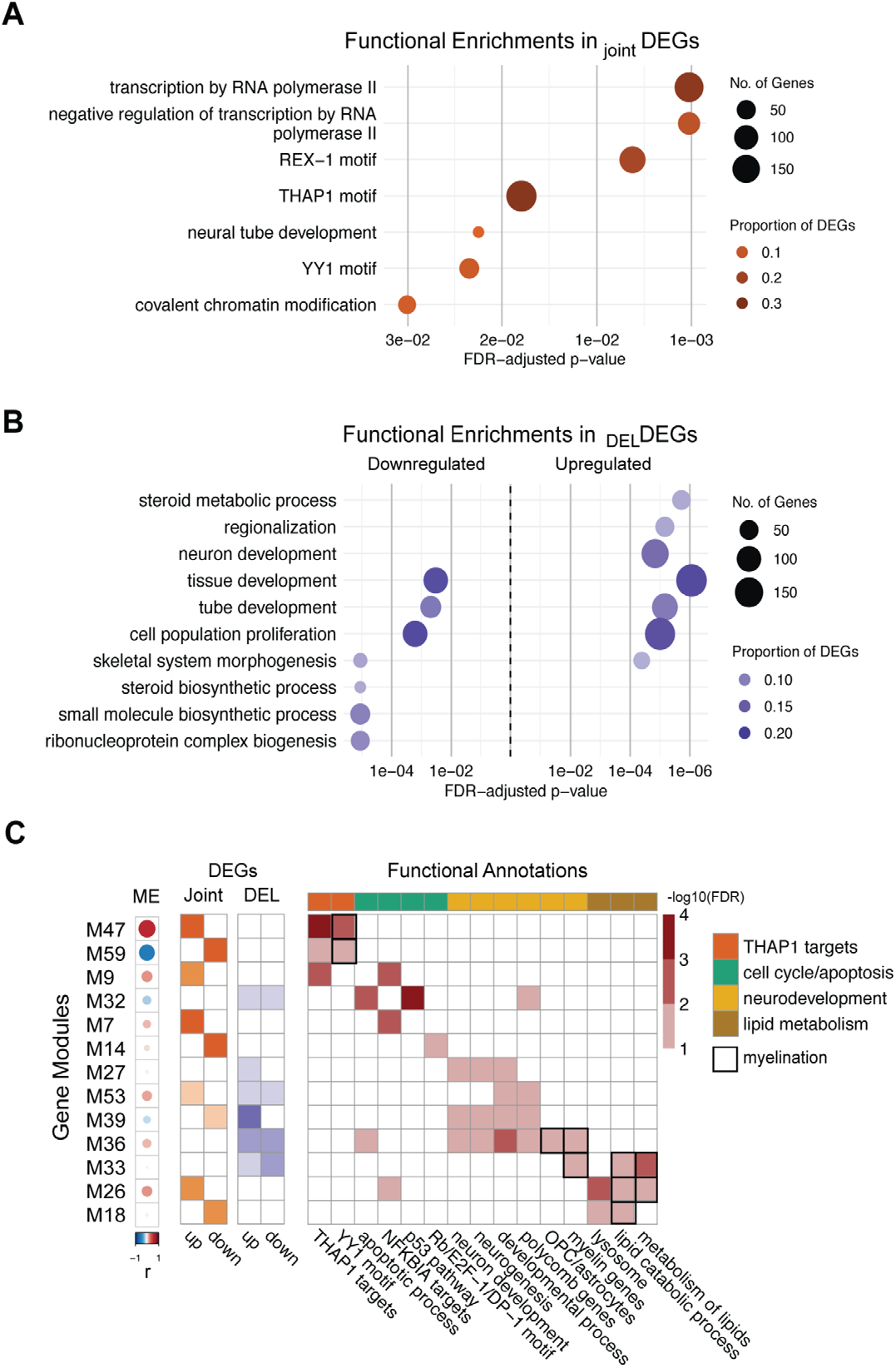
*Functional analysis reveals convergent signatures of neurodevelopment and myelination across diverse* THAP1 *mutations*. **(A)** Functional analysis via GO term enrichment testing in DEGs identified transcription related terms in _JOINT_DEGs. Additionally, “neural tube development” was an enriched GO term. As expected, the THAP1 binding motif was overrepresented in these DEGs, given that the mutations that comprise the meta-analysis functionally alter THAP1 binding to its targets. YY1, which co-binds THAP1-bound sequences and is a known regulator of oligodendrocyte development, is also overrepresented, as well as REX-1, a transcription factor that has known activity in stem cells. All terms plotted have FDR < 0.05 in Fisher’s tests for enrichment. **(B)** _DEL_DEGs were enriched for GO terms related to development, including specifically, “neuron development” (FDR 1.5×10^-5^), as well as “cell population proliferation,” consistent with known functions of THAP1. _DEL_DEGs were not enriched for THAP1 targets. Additionally, _DEL_DEGs were enriched for GO terms related to lipid metabolism or synthesis (in both up/down-regulated DEGs, FDR < 1.0×10^-5^ in both). All terms plotted, FDR < 0.01 in Fisher’s tests for enrichment. In both of these enrichment plots (A and B), circle size represents the number of intersecting genes between each gene set (y-axis) and DEGs, while color/shade represents the proportion of DEGs harboring a certain gene set term. For visualization, GO terms were reduced from the entire list of enriched terms using semantic similarity measures, as described in the Methods. **(C)** Modules of co-expressed genes where the module eigengene (ME) correlates/ anticorrelates with THAP1 expression (r, biweight mid-correlation) were probed for enrichment of _JOINT_DEGs and _DEL_DEGs (all shaded boxes corresponding to DEGs with p-value < 0.01 in Fisher’s test for enrichment). Modules were then annotated using GO terms and custom lists, revealing significant enrichment of terms that are related to known THAP1 functions in humans and mice (cell cycle control, apoptosis, and neural differentiation). This analysis also revealed modules that were associated with lipid metabolism and lysosomal function, as well as overlapping modules that are related to myelination. The module that is most correlated with *THAP1* expression (i.e., M47) includes *THAP1* itself, is enriched for the THAP1 motif (FDR 0.01), and for the YY1 motif (FDR 0.02), which include genes known to participate in oligodendrocyte development and differentiation, a critical pathway in myelin formation. Enrichment of annotations FDR < 0.10 when shaded in the heatmap using Fisher’s test for enrichment; annotations/terms were curated for visualization, and each are defined in the Methods.

Although significant _JOINT_DEGs and _DEL_DEGs had minimal overlap, gene annotations of _JOINT_DEGs were also enriched for terms related to neuron development (e.g., neural tube development FDR 0.02, ***Figure 3A***) and overrepresented with genes exhibiting selective pressure and loss-of-function intolerance (FDR 0.01, using genes with upper bound of the observed/expected confidence interval < 0.25 in gnomAD) that are enriched in neurodevelopmental disease.^45,60,61^ Further, query for overrepresented transcription factor binding motifs in _JOINT_DEGs identified not only the THAP1 motif, but also elements for YY1 and REX-1 (both FDR < 0.05, ***Figure 3A***), both of which have roles in neurodevelopment.^62–65^ Taken together, functional enrichment analyses suggest that convergent pathways and networks related to development characterized both _JOINT_DEGs and _DEL_DEGs, and imply neurodevelopmental consequences resulting from diverse classes of *THAP1* mutations and across different levels of *THAP1* dysregulation. This effect was more pronounced in haploinsufficiency, as previously shown.

To further probe the functional impact of *THAP1* perturbation, we performed network-based analyses to identify the modules of co-regulated genes representing active biological processes in our data set.^66^ We correlated module eigengenes (MEs) with *THAP1* expression, then layered with significant overlap with _JOINT_DEGs and _DEL_DEGs, to select the gene modules that were impacted by our mutational modeling. The two modules with MEs most correlated with *THAP1* expression (module 47 [biweight midcorrelation r = 0.58; FDR 1.4×10^-7^,] and module 59 [r = −0.50; FDR 1.7×10^-5^]) contained the highest number of _JOINT_DEGs and were strongly enriched for THAP1 DNA targets, as expected. Overall, DEGs were distributed across 13 *THAP1*-associated modules that were not necessarily enriched with known THAP1 targets, suggesting that the transcriptional effects captured by our profiling and analyses were not exclusive to direct regulation by THAP1 but may also include downstream and secondary effects. Annotation of gene modules revealed functions that have previously been described for THAP1 in human and/or mouse cells, such as apoptosis and cell-cycle control via the p53 pathway and the pRB/E2F transcription factor, and neurodevelopmental processes (***Figure 3C***),^13,58,59,67^ showing that our workflow and analyses are able to recover expected effects based on known *THAP1* biology. Expanding and recapitulating the results of enrichment analyses performed on DEGs, we also discovered *THAP1*-associated gene modules that were functionally related to lipid metabolism, lysosomal function, glial development (i.e., oligodendrocyte progenitor cells [OPC] and astrocytes), and myelination (all FDR < 0.10). Module 47 in particular included *THAP1*, was overrepresented with _JOINT_DEGs (38% of the genes in the module overlapped with _JOINT_DEGs, Fisher’s test p-value < 2.2×10^-16^) and was enriched for YY1 (FDR 3.5×10^-7^), a transcription factor with critical functions in oligodendrocyte differentiation^63^.

### *Thap1* disruption is associated with defects in myelination in mouse models

The transcriptional profiling of NSCs thus indicated that *THAP1* disruption is associated with gene modules related to myelination. To directly assess the effects of *THAP1* perturbation on myelin gene expression and morphology, we analyzed two *Thap1* mouse models harboring: (1) knock-in of the C54Y missense mutation (C54Y-KI, *Thap1*^C54Y/+^) that we profiled in iPSCs, and (2) a deletion allele based on knock-out of *Thap1* exon 2 (ex2-KO, *Thap1*^-/+^). Both mice display subtle motor abnormalities and impaired performance in behavioral assays compared to WT littermates.^68^ We interrogated three brain regions that have been implicated in the pathogenesis of dystonia, including regions where microstructural white matter abnormalities have been documented via neuroimaging studies in humans,^69,70^ and observed age-, region-, and genotype-specific variability in gene expression compared to control tissue. In the cerebellum and cortex, we detected increased expression of myelin-related genes (*Cnp*, *Mag*, *Mobp*, *Mog*, and *Plp1*) in postnatal day 21 (P21) mice, which was most significant in the ex2-KO model (n = 5 C54Y-KI, 7 WT littermates; n = 10 ex2-KO, 12 WT littermates). In 2-month-old (adult) mice, we detected a consistent and significant downregulation of these genes in the cortex (p-value < 0.05, n = 4 C54Y-KI, 4 WT mice, ***Figure 4A***). There was also a significant reduction in myelin structural integrity based on LFB staining in the anterior commissure of C54Y-KI and ex2-KO mice (p-value < 0.05, ***Figure 4B***) and in the corpus callosum of ex2-KO mice (p-value < 0.01; n = 4 C54Y-KI, 5 WT mice; n = 8 ex2-KO, 8 WT mice, ***Figure 4C***). These data appear consistent with the transcriptional analyses and indicate that dysregulation of *Thap1* activity impacts the formation and/or maintenance of myelin in the brain.

**Figure 4.**
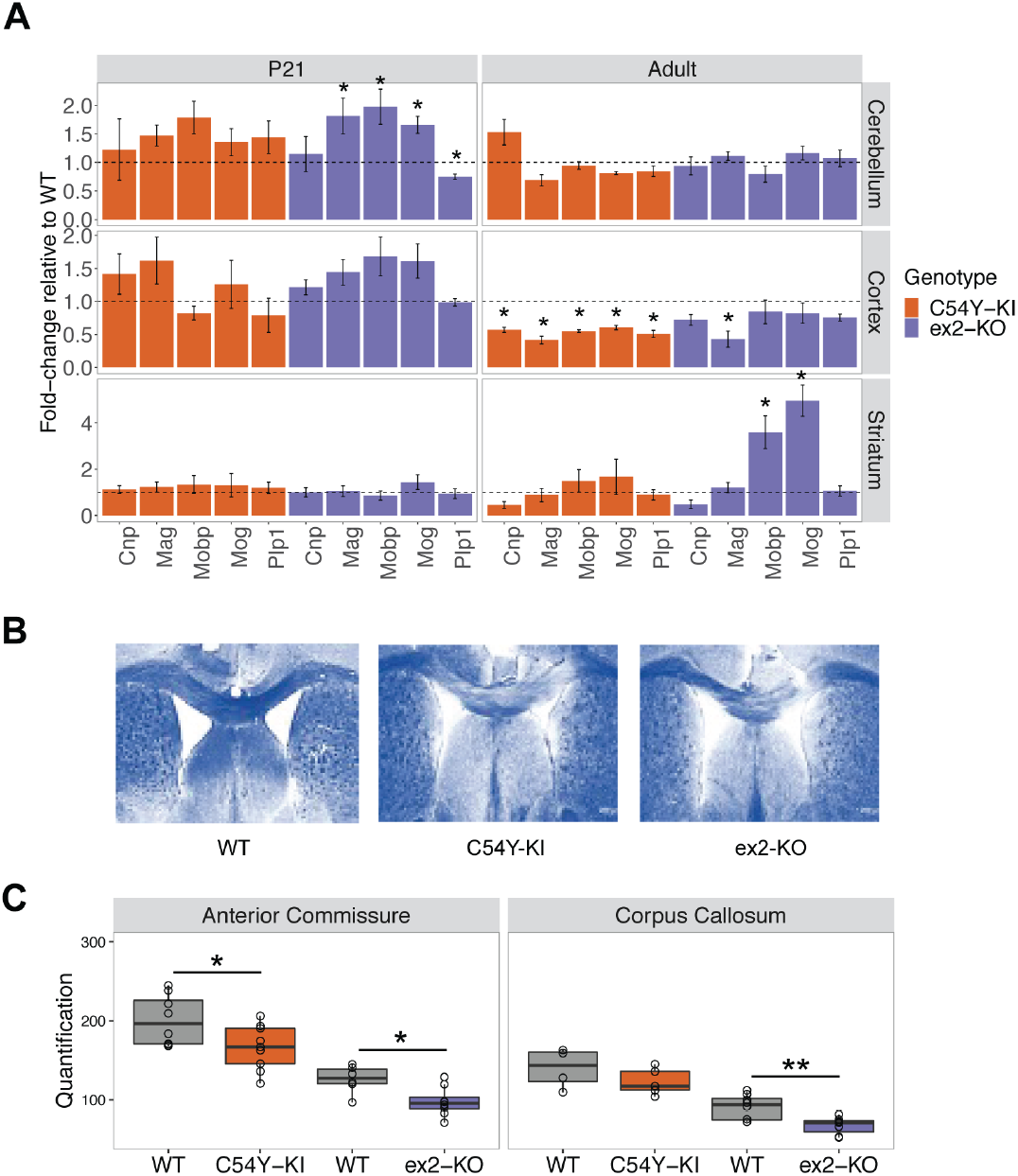
Thap1 *disruption is associated with defects in myelination in mouse models*. **(A)** Quantitative PCR assays revealed significant decreases in the expression of myelin-related genes in the cortex of 2-month old mice (adult) with heterozygous knock-in of *Thap1* C54Y (C54Y-KI, *Thap1*^C54Y/+^) and deletion of exon 2 (ex2-KO, *Thap1*^-/+^) compared to WT littermates. These reductions were not seen in younger mice (postnatal day 21 [P21]). Further, significant increases in gene expression in the cerebellum of P21 mice as well as in the striatum of adult mice with ex2-KO was seen. For P21, n = 5 C54Y-KI, 7 WT littermates; n = 10 ex2-KO, 12 WT littermates; for adult, n = 4 mice per group. Quantification is shown as FC against mean expression in WT littermates. Error bars represent standard error of the mean, and significance was tested via Student’s t-tests on normalized expression values in mutant vs. WT groups: * p-value < 0.05. See also ***Supplementary Figure 5***. **(B)** Representative LFB staining of myelin structural integrity in the corpus callosum of adult WT, C54Y-KI, and ex2-KO mice, showing myelin loss in adult mice with mutant *Thap1* alleles. **(C)** Quantification of LFB stain intensity in 25 adult mice (n = 4 C54Y-KI, 5 WT; n = 8 ex2-KO, 8 WT) revealed significant myelin loss in the anterior commissure of C54Y-KI and ex2-KO mice and in the corpus callosum of ex2-KO mice. Student’s t-test on stain quantification: * p-value < 0.05, ** p-value < 0.01.

## DISCUSSION

In recent years, advances in human genetics and genomics have led to the discovery of an expanding list of genetic variation associated with hereditary dystonia. This list currently includes genes such as *TOR1A*, in which a single mutation has been linked to dystonia in multiple populations^71^ versus others, such as *THAP1*, *KMT2B*, *ANO3*, *GNAL*, *TAF1* and *ATP1A3*, in which varied alterations have been associated with disease.^6,40^ In some of these genes harboring multiple mutations (e.g., *THAP1*, *GNAL*, *ANO3*), genetic variation results in a largely consistent neurologic phenotype, whereas in others, such as *ATP1A3* and *TAF1*, diverse mutations result in distinct clinical syndromes.^72–75^ Dissecting the complex relationships between genetic variation and human disease requires a better understanding of their functional outcomes, the ways in which they differ, and, in particular, the affected cellular pathways that they share.

Previous efforts to define such convergent mechanisms have gained insight from studies of: (1) the impact of dystonia-specific genetic variation within structural domains of a target protein^76^; (2) functional inter-relationships among proteins implicated in dystonia^4,77^; and (3) overlapping brain expression patterns of genes associated with dystonia.^78^ Here we have outlined an additional approach that leverages an allelic series of dystonia-specific mutations in near-isogenic iPSC-based neuronal models to infer mutational mechanisms from their transcriptome-wide effects. By engineering sequence changes into an isogenic background, we isolated the effects of each mutation while capturing variation through the analysis of multiple clones per genotype. In the differential expression analyses, each mutational group was compared internally to its own set of controls, enabling systematic and repeated determinations of dysregulated genes. In this context, the overlap of differentially expressed genes across multiple sets, with each independent analysis meeting stringent criteria for statistical significance, was considerably beyond what could be expected by chance, and thus defined a high-confidence signature of *THAP1* disruption. We used meta-analytic methods to expand this core set of dysregulated genes, after obtaining proof of highly correlated mutational expression profiles. This experimental and analytical pipeline mitigates current challenges in discovery using functional genomics in iPSC-based models, i.e., noise introduced by individual genetic variation is resolved by the isogenic background, while false positive signals resulting from off-target genome-editing effects are reduced in the meta-analysis of results across an allelic series. Lastly, we performed validation in mice, which recapitulated the discovered transcriptomic signatures and implied physiologically relevant effects in a model organism and in different developmental time points.

The pathogenic missense mutants analyzed here have behaved differently in previous biochemical assays, raising the possibility that they might exert unique transcriptional effects. Unlike the S21T and P26R variants, the N12K mutant did not affect DNA-binding affinity but rather decreased thermostability.^14,17^ The C54Y mutation may also be unstable, as the substitution targets one of four residues that together bind a single zinc ion that may be required for structural integrity.^14,15^ In our modeling, we found that the global transcriptional effects of these mutations are highly similar, implying a common perturbed pathway related to the regulatory effects of *THAP1* that is independent of the mutation’s impact on THAP1 stability. In comparing the signatures across the allelic series, we observed a stepwise increase in transcriptional effects induced by the heterozygous, homozygous, and haploinsufficiency models, with the latter inducing dysregulated genes that was two orders of magnitude greater than that of the missense mutations. Although we hypothesized that the R146 frameshift variant would be the functional equivalent to the deletion if the loss of residues in the NLS prevented nuclear import, its signature correlated with those of the missense mutants in the DNA-binding domain. These results imply two competing mutational mechanisms. One possibility is that variation in the DNA-binding domain and NLS represent hypomorphic alleles, consistent with previous studies in mice reporting that exon deletion produced broader effects than heterozygous knock-in of the C54Y allele^16,67^ and that C54Y fails to rescue embryonic lethality of *Thap1*-null.^59,68^ Alternatively, missense variation in the DNA-binding domain may induce a novel gain-of-function in the cytoplasm, with that property being responsible for some of the shared expression changes with the R146 mutation that abolishes nuclear import of THAP1. To resolve these scenarios, future studies will be required to identify cytosolic functions of THAP1 and to determine if the R146 and similar mutations in this domain retain some capacity for passive nuclear entry and DNA binding. As predicted, there was a paucity of transcriptional effects induced by the M143V variant suggesting it could be benign. However, given the position of this mutation within the coiled-coil domain, we cannot exclude the possibility that it may affect critical interactions with partner proteins that do not directly disrupt transcriptional activity.

Although the dysregulated genes associated with missense versus haploinsufficiency models were largely exclusive, they appeared to converge on common cellular pathways associated with neurodevelopment, lipid metabolism, and myelin. We identified dysregulation of myelin-related transcripts associated with *THAP1/Thap1* disruption, including consistent downregulation of galactosylceramidase (*GALC*), which encodes a lysosomal enzyme that catabolizes the major lipid constituents of myelin. Mutations in the gene cause an autosomal recessive neurodevelopmental disorder with leukoencephalopathy and demyelination,^79^ and enzyme deficiency in mice results in reduced proliferative potential in NSCs and functional impairment of neuronal and oligodendroglial progeny.^80^ Further, we identified modules of genes that were correlated with *THAP1* perturbation and functionally associated with YY1 targets, lipid metabolism, and myelination. Previous studies in mice have linked *Thap1*-haploinsufficiency to impaired maturation of oligodendrocytes and loss of YY1 binding to a network of genes related to oligodendroglial differentiation.^59^ Our results extend these observations further to human disease-specific variation and suggest that a complex interplay of dysregulated lysosomal sphingolipid metabolism, disrupted oligodendrocyte differentiation, and altered myelin formation and/or maintenance may contribute to the molecular and cellular mechanisms underlying *THAP1* disease.^81–84^ Although overt demyelination has not been specifically documented in dystonia, neuroimaging studies have reported white matter abnormalities in multiple forms of the disease, including: (1) syndromes associated with genetic variation in *TOR1A*, *THAP1*, *SGCE*, *TAF1*, *KMT2B*, and *COL6A3*^20,69,85–89^; and (2) idiopathic cases of cervical dystonia, writer’s cramp, laryngeal dystonia, and blepharospasm.^90–93^ The mechanisms underlying these disturbances in white matter tracts are not known, and further investigation is needed to determine if they reflect common defects in any of the pathways perturbed by genetic variation in *THAP1*.

Identifying cellular deficits that may be shared among multiple forms of dystonia remains a high priority for translational research in these disorders. Monogenic dystonias are rare syndromes, and if convergent disease mechanisms exist, they may offer opportunities for broad therapeutic intervention. In this study we outlined a systematic experimental and analytical variant-to-function platform of iPSC-based modeling and genome editing that we initially used to understand the cellular consequences of genetic variation in *THAP1*. These tools may similarly facilitate future studies that seek to understand how diverse mutations across multiple genes result in similar phenotypes, while enabling the potential discovery of common disease targets.

## ACKNOWLEDGMENTS

Funding for these studies was provided by National Institutes of Health grants NS102423 (MET, DCB, LJO) and NS087997 (DCB, LJO, NS). AD was a recipient of fellowships from the Dystonia Medical Research Foundation and the MGH Collaborative Center for X-linked Dystonia Parkinsonism.

## SUPPLEMENTARY FIGURES

**Supplementary Figure 1.**
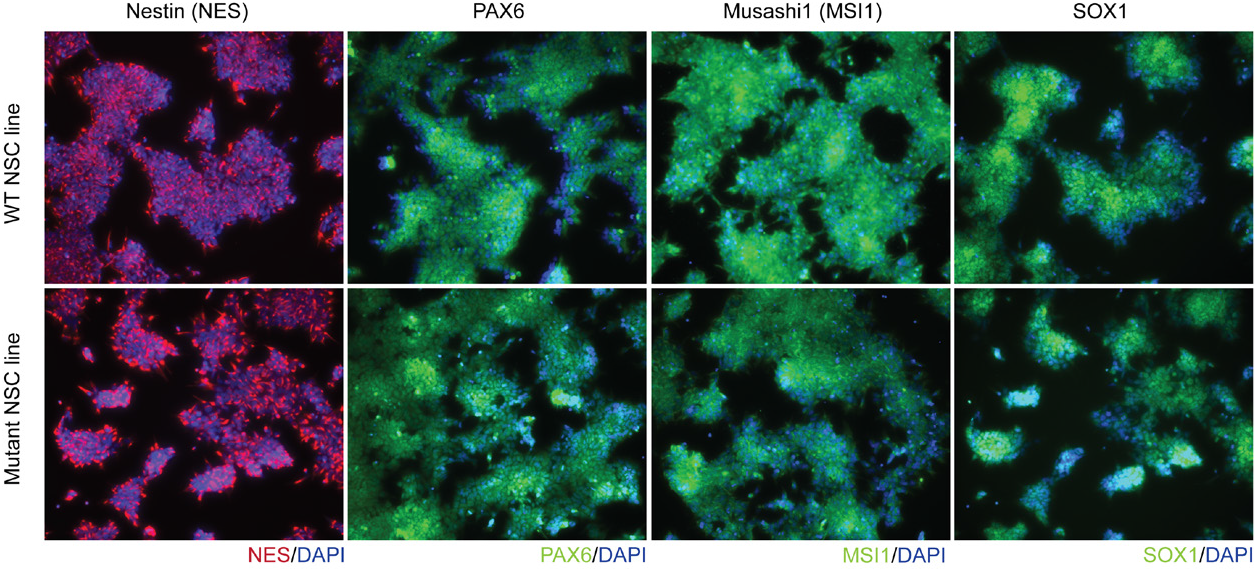
Related to ***Figure 1B***. Immunocytochemistry performed using antibodies against known markers of the NSC lineage. This representative subset of images shows that there was identical expression of NES, PAX6, MSI1, and SOX1 in mutation and WT NSC lines. There were likewise no obvious growth nor morphological differences observed between genotypes, and across differentiation batches.

**Supplementary Figure 2.**
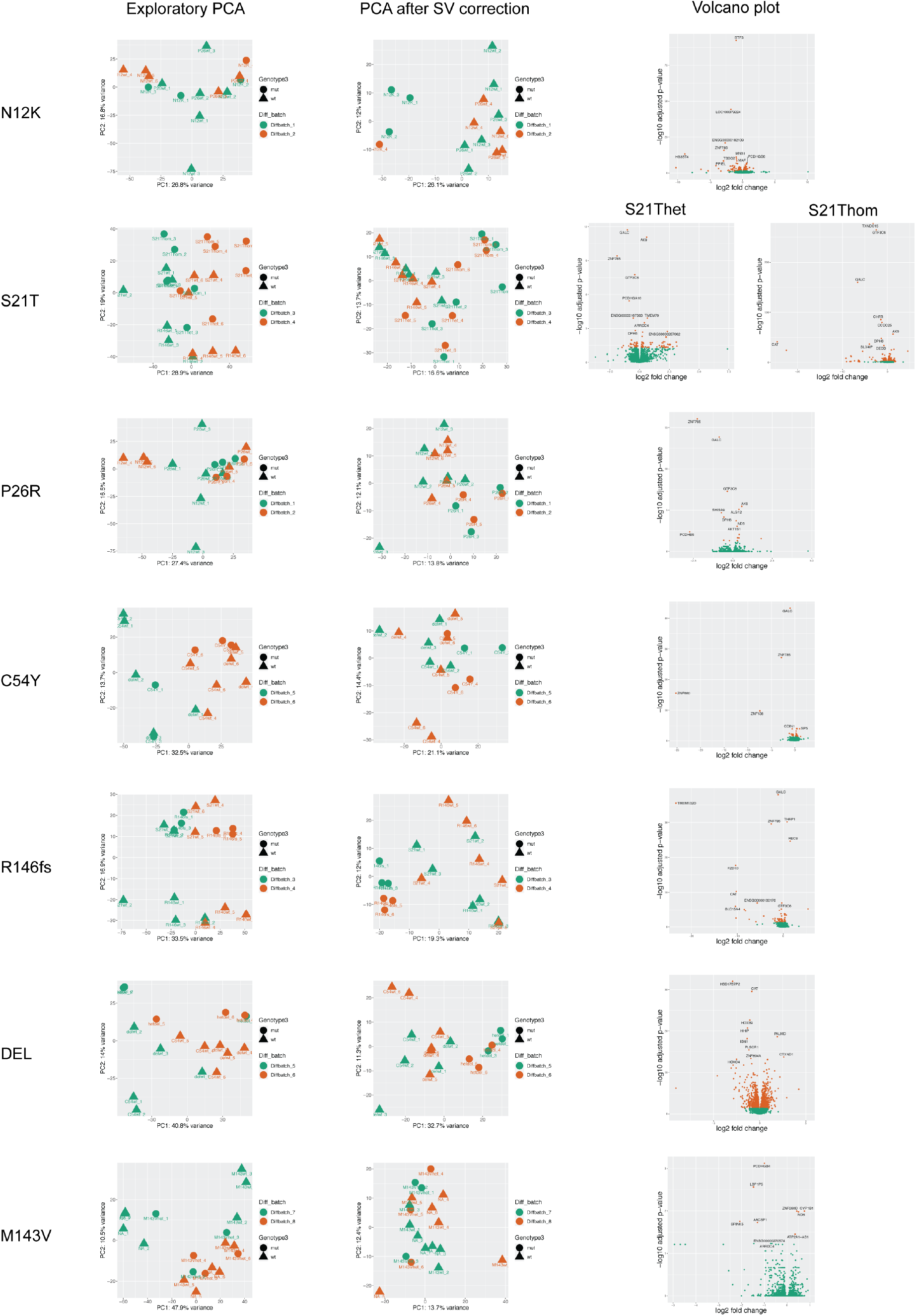
Related to ***Figure 2B***. Exploratory/uncorrected PCAs revealed that differentiation batch effects dominated the variance in expression profiles. These were however ameliorated via SV correction, resulting in PCAs where profiles clustered according genotype, in accordance with the intended analysis model. DEGs were discovered at FDR < 0.05 in comparisons of single-nucleotide (i.e., N12K, S21Thet, P26R, C54Y) or small deletion (i.e., R146fs) mutations vs. corresponding controls. The largest transcriptomic effect in terms of number of DEGs was seen in DEL. In the volcano plots, orange points are genes with FDR < 0.05 in the differential expression analyses, and the genes with the lowest adjusted p-values are labeled.

**Supplementary Figure 3.**
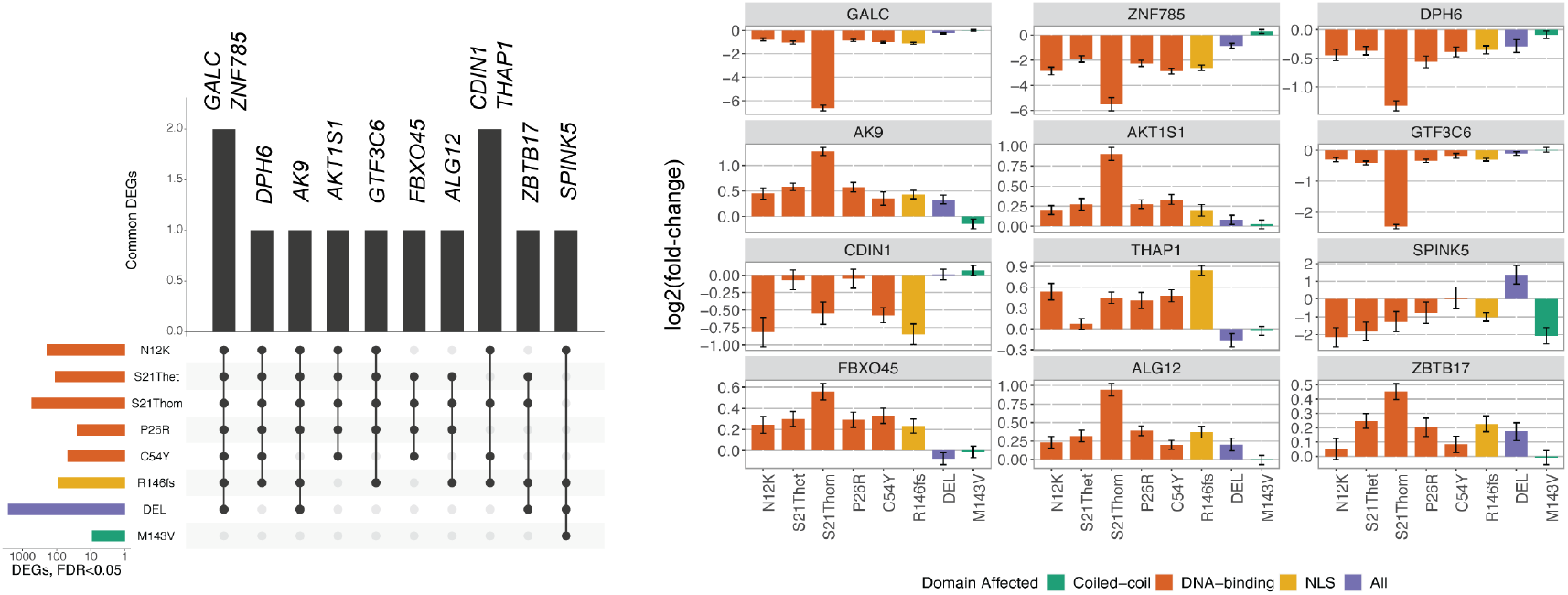
Related to ***Figure 2B***. Overlapping DEGs (FDR < 0.05) across different comparisons (i.e., exclusive sets of mutant and control samples for each comparison). Analysis of expression patterns of overlapping DEGs across different mutations in the allelic series revealed consistently downregulated genes across all pathogenic mutations and DEL: *GALC*, *ZNF785*, *GTF3C6*, and *DPH6*, while consistently upregulated genes include *AK9*, *AKT1S1*, *ALG12*, and *ZBTB17. THAP1* is upregulated by mutations in the DNA-binding domain/NLS, while it is downregulated in the DEL model. Error bars represent standard error of the log2FC.

**Supplementary Figure 4.**
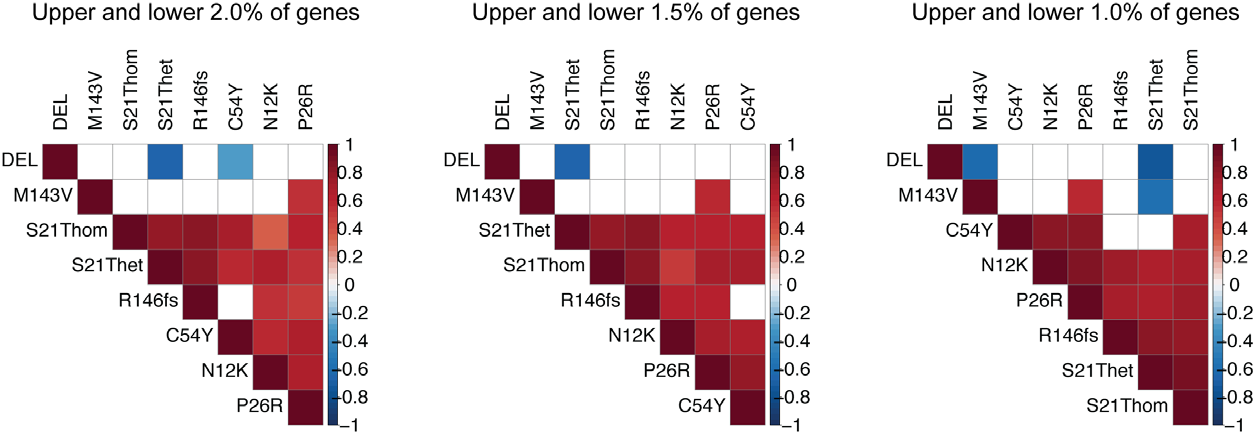
Related to ***Figure 2C***. Mutation-pairwise rank-order (q-value) correlations across different q-value thresholds representing the most differentially expressed genes (4%, 3%, 2% of all genes profiled) robustly identify overlapping signatures induced by the N12K, S21T, P26R, C54Y, and R146fs mutations, that is exclusive to the signature of DEL and not consistently identified in M143V model (i.e., benign variant). Spearman’s correlations (r): p-value < 0.05.

**Supplementary Figure 5.**
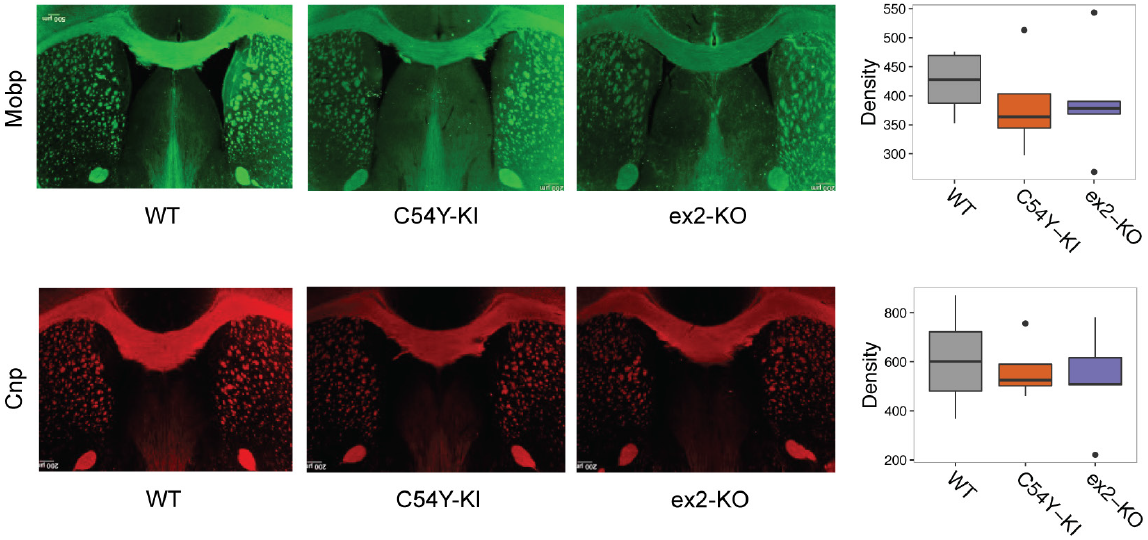
Immunocytochemistry (representative image and quantification in 16 mice) performed on the corpus callosum revealed non-significant reductions of mean intensity density for both adult C54Y-KI and ex2-KO mouse compared to WT, for two myelin-related proteins, Mobp and Cnp (mean intensity densities: Mobp WT = 423.4, C54Y-KI = 384.4, n.s., ex2-KO = 389.7, n.s.; Cnp WT = 606.8, C54Y-KI = 566.7, n.s., ex2-KO = 526.6, n.s.; n = 4 C54Y-KI, 5 ex2-KO, 7 WT mice). Student’s t-test for group means: all p-values ≥ 0.05 (n.s.).

## SUPPLEMENTARY TABLES

The following Supplementary Tables are available from the authors upon request.

Supplementary Table 1. Metadata, summary of quality control metrics, and results of differential expression analyses in different groups of mutant vs. control across the allelic series.

Supplementary Table 2. Off-target effects queried and identified in different mutation groups.

Supplementary Table 3. Results of weighted Z-score-based meta-analysis of DEGs in the N12K, S21Thet, P26R, C54Y, and R146fs groups (full and sign-test applied).

Supplementary Table 4. Results of GO term and transcription factor binding motif enrichment analysis in _JOINT_DEGs and _DEL_DEGs (full lists and reduced lists based on semantic similarity). Supplementary Table 5. Results of WGCNA, including module membership and overlaps with differentially expressed genes. Functional annotation of modules of interest using GO term and transcription factor binding motif enrichment analysis.

## Notes

### Competing Interest Statement

The authors have declared no competing interest.

